# Experimental reproduction numbers disentangle vaccine effects on susceptibility and infectiousness during H5N1 transmission in geese

**DOI:** 10.64898/2026.09.02.748824

**Authors:** Ronja Piesche, Jana Schulz, Michael Höhle, Martin Beer, Christian Grund, Timm Harder, Jose L Gonzales

**Author notes:** Corresponding authors: Timm Harder and Christian Grund. These authors contributed equally to this work (Ronja Piesche, Jana Schulz).

## Abstract

Vaccination against high pathogenicity avian influenza virus (HPAIV) is increasingly used to protect poultry, but vaccine performance is commonly inferred from clinical protection and virus shedding rather than measured transmission. We asked whether reproduction numbers from controlled transmission experiments can quantify how vaccination changes infection dynamics in virus-exposed geese, i.e. susceptibility and infectiousness.

Domestic geese were prime-boost vaccinated with an H5 clade 2.3.4.4b RNA-amplicon vaccine and challenged with homologous HPAIV H5N1. Replicats of seeder-sentinel groups represented transmission among unvaccinated animals, to vaccinated contacts, and from vaccinated breakthrough-infected animals. Vaccinated directly challenged geese remained clinically protected although all became productively infected. Estimated reproduction numbers were R_0_=4.7 (95% CI, 2.8-8.0) among unvaccinated geese, R_s_=2.6 (1.6-4.5) for transmission to vaccinated contacts, R_i_=3.7 (2.0-6.8) for transmission from vaccinated infected geese, and R_vacc_=2.0 (0.8-4.9) for a fully vaccinated population.

Vaccination reduced transmission but, under these experimental exposure conditions, did not reduce the point estimate for R_vacc_ below one. Even though vaccinated infected geese shed substantially less viral RNA, infectiousness was sufficient to sustain transmission in vaccinated geese, indicating that RNA shedding alone may not reliably predict efficacy of transmission reduction. Experimental reproduction numbers therefore provide a direct population-level complement to conventional vaccine testing protocols and can separate effects on susceptibility from effects on onward transmission.

**Author Summary:** Vaccines against high pathogenicity avian influenza are usually evaluated by asking whether they prevent disease and reduce the amount of virus shed by infected birds. For disease control, however, it is equally important to know whether vaccination prevents onward transmission. We used domestic geese infected with H5N1 avian influenza virus to test whether this effect can be measured directly in small transmission experiments. By co-housing vaccinated and unvaccinated infected birds with vaccinated or unvaccinated contact birds, we estimated reproduction numbers for different transmission circumstances. Vaccination fully protected geese from clinical disease and reduced viral RNA shedding, but it did not prevent infection or transmission. The estimated reproduction number fell from 4.7 among unvaccinated geese to 2.0 for transmission to a fully vaccinated population, although the confidence interval was wide. Notably, the reduction in viral RNA shedding overestimated the effect on infectiousness. Thus, shedding alone may overstate how strongly a vaccine limits transmission. Directly estimating transmission can therefore add important information to conventional vaccine-efficacy studies and may improve comparisons between vaccines.

## Introduction

High pathogenicity avian influenza virus (HPAIV) H5N1 of clade 2.3.4.4b has established a panzootic that affects poultry and wild birds on a global scale and is accompanied by recurrent spillover into mammals, including humans [1,2]. Outbreak control relies on rapid detection, movement restrictions, culling of affected flocks, and strict biosecurity, but sustained circulation in wild-bird reservoirs makes repeated incursions into poultry difficult to prevent completely [3]. Consequently, preventive vaccination is increasingly being considered or implemented as an additional control measure [4–6].

For vaccination to contribute to population-level control, protection of individual animals from clinical disease is not sufficient. Vaccination must also reduce the probability that exposed animals become infected and/or reduce infectiousness of breakthrough-infected animals, thereby limiting onward virus transmission. Veterinary vaccine studies often deduced these effects from surrogate endpoints, particularly reductions in viral shedding, and sometimes from infection of a limited number of sentinel animals [7,8]. Although these measures are informative, they do not directly quantify the combined transmission process and may be misleading if the relationship between molecular shedding and infectiousness differs between vaccinated and unvaccinated hosts.

Transmission can be summarized by the reproduction number R, which integrates the rate at which infectious hosts generate new infections and the duration for which they remain infectious [9,10]. In a fully susceptible population, the basic reproduction number R_0_ provides a baseline for transmission. Vaccination can then be considered as acting through two biologically distinct components: reduced susceptibility of vaccinated recipients and reduced infectiousness of vaccinated breakthrough cases. Estimating these components separately is particularly relevant for HPAIV because vaccines may prevent severe disease while still allowing infection and virus replication.

Small-scale seeder-sentinel experiments provide a tractable setting in which these transmission components can be estimated under controlled exposure conditions. Their advantage is not that the resulting reproduction numbers reproduce field values, but that variation in host composition, contact structure, and exposure can be constrained sufficiently to compare transmission pathways and vaccine effects. A central challenge is secondary transmission from newly infected sentinels, especially when susceptible contacts rapidly become infectious. Analytical approaches therefore are considered to require a distinction of transmission attributed only to inoculated seeders from transmission generated by all infectious animals present in a group.

Here, using domestic geese and homologous HPAIV H5N1 challenge, we estimated R_0_ for transmission among unvaccinated animals, R_s_ for transmission from unvaccinated infected animals to vaccinated recipients, and R_i_ for transmission from vaccinated infected animals to unvaccinated recipients. These quantities were used to derive R_vacc_ for a fully vaccinated population and to decompose vaccine efficacy into effects on susceptibility and infectiousness. We tested the robustness of the model for effects of secondary transmission from newly infected sentinels by comparing estimates from models in which transmission was restricted either to inoculated seeders only or which also allowed secondary transmission from infected sentinels. We further asked whether reductions in viral RNA shedding tracked the reduction in infectiousness measured from observed transmission.

## Results

For collection of comparable data sets to analyze transmission and quantify R, 24 prime-boost vaccinated geese and 48 unvaccinated geese were subjected to an HPAIV challenge infection following assignment to three clusters (I-III). Each cluster was subdivided into four subclusters of six animals each, comprising three inoculated seeder animals and three sentinel animals, as illustrated in Figure 1. Geese were prime-boost vaccinated using a commercial H5 clade 2.3.4.4b RNA-amplicon vaccine Before challenge infection,) the vaccinated animals, with few exceptions, tested seropositive by AIV H5 ELISA (according to the manufacturer’s cut-off criteria), and all showed HI titers against the homologous H5N1 clade 2.3.4.4b antigen with a mean log₂ titer of 5.9 ± 1.2 (range, 4–8) (see Supplementary Table 1). They remained seronegative by AIV NP ELISA until challenge, confirming that antibodies were due to vaccination, ruling out any contact to replication-competent AIV. All unvaccinated geese remained serologically negative in all assays (Supplemental table 1). The three cluster configurations were used to distinguish baseline transmission, vaccine effects on susceptibility, and vaccine effects on infectiousness; detailed experimental and statistical procedures are provided in Materials and Methods.

**Figure 1.**
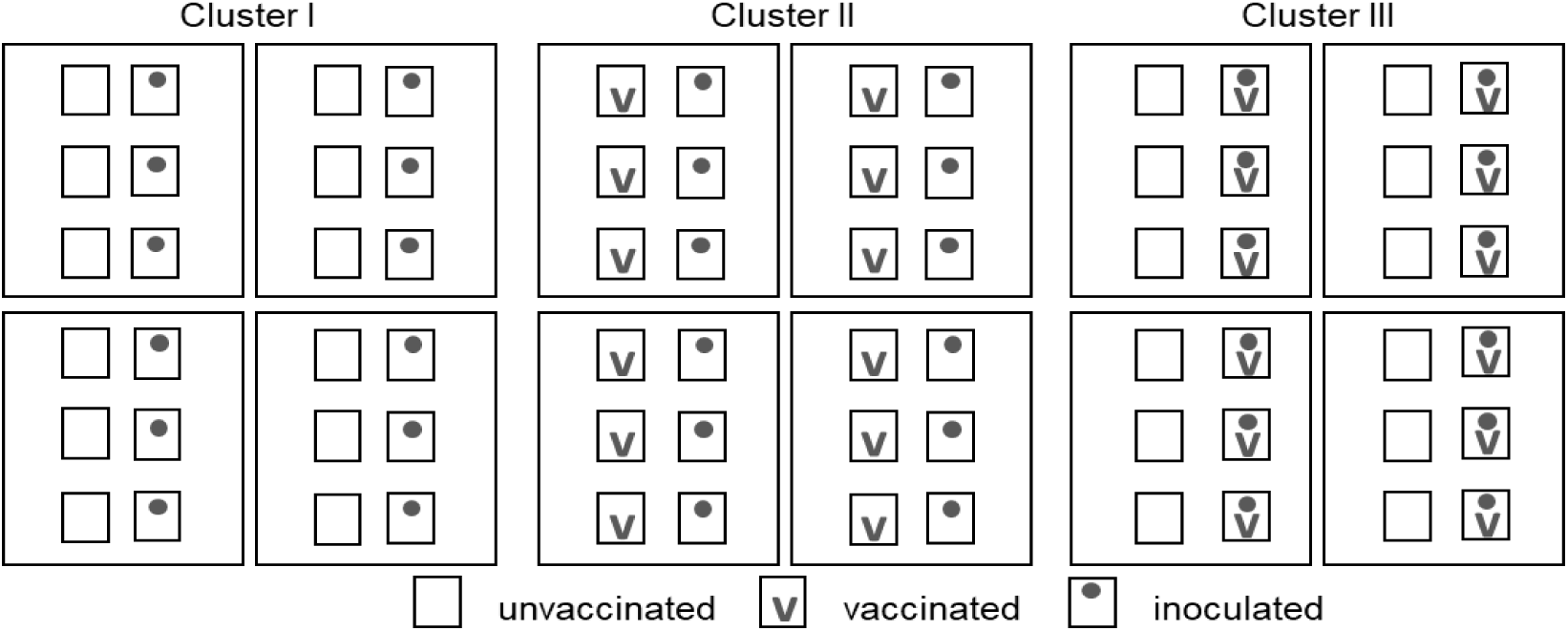
Generalized layout of animal experimental design. Essentially, test subjects were randomly assigned to three population clusters each of 24 individuals (total, n=72 geese). Within each cluster, four subclusters of six individuals were formed. In cluster I, all six individual test subjects of a subcluster have the same immunological status (“susceptible”); in other clusters each three individuals of a subpopulation share the same immune status with respect to pathogen-specific immunity, i.e. vaccinated (‘v’ squares) or not (blank squares). Each three individuals (‛seeders’) of a subpopulation are inoculated with the pathogen under study, here: HPAIV H5N1 (dotted squares). Individuals not inoculated (‛sentinels’) are co-housed with the inoculated ones 16 hours after inoculation.

### Clinical protection does not prevent infection after direct H5N1 challenge

#### Vaccinated seeders become infected with similar timing but remain clinically protected

Viral RNA in oropharyngeal swabs was generally used to define infection and the PCR-based infectious period because detection occurred earlier and at higher genome-equivalent concentrations than in cloacal swabs (Supplementary Table 2). Seeder and sentinel birds were co-housed from 16 hours post inoculation (hpi), defined as 0 hours post contact (hpc). In cluster I, eight of twelve unvaccinated seeders were RT-qPCR-positive at 16 hpi, two additional birds at 24 hpi, and the remaining two at 40 hpi, with swab loads ranging from 4.47 x 10^2 to 7.37 x 10^6 Geq/ml. Four seeders developed clinical disease by 64 hpi and three additional birds by 88 hpi; all seven reached predefined humane endpoints within 24 hours after the first clinical signs. The remaining five seeders were euthanized after all cluster-I sentinels had become RT-qPCR-positive, before clinical signs developed. In cluster II, 10 of 12 unvaccinated seeders were positive for viral RNA by 16 hpi and the remaining two by 24 and 40 hpi, respectevely. Eleven animals were euthanized at 96 hpi, seven after acute neurological signs, and the final bird was removed at 120 hpi after all sentinels had become positive. Vaccinated inoculated seeders in cluster III also became RT-qPCR-positive within 48 hpi (16 hpi, n=10; 24 hpi, n=1; 48 hpi, n=1), but none developed clinical signs during 14 days of observation. Survival regression showed no significant difference in time to PCR-defined infectiousness (“latent period”) between vaccinated and unvaccinated seeders (p=0.14); the overall median was 9.7 h (95% CI, 2.6-36) (Figure 2).

**Figure 2.**
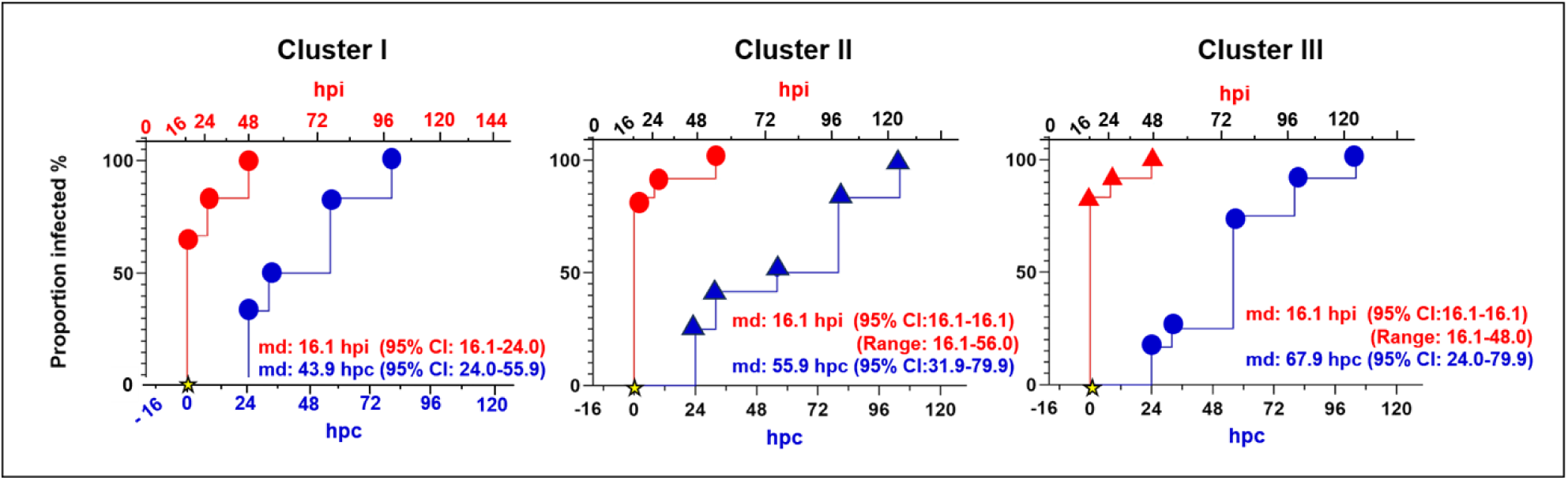
Kaplan–Meier curves showing infection dynamics for seeder and sentinel geese in clusters of experimental transmissions. Each panel represents one cluster and compares two groups: inoculated seeder animals (red) and sentinel animals (blue). The plots display two synchronized x-axes: the upper red axis shows hours post inoculation (hpi) for inoculated seeder geese, and the lower blue axis shows hours post contact (hpc) for sentinel geese. The start of co-housing (contact time = 0) is marked with a star on the lower x-axis. In all clusters, unvaccinated animals are shown as circles, while in clusters II and III vaccinated geese are shown as triangles. The y-axis represents the proportion of infected animals (%) as detected by RT-qPCR in oropharyngeal swabs. The numbers shown in red and blue indicate the median time until start of the infectious period for inoculated seeder (red) and sentinel (blue) geese.

Transmission to sentinels occurred in all three clusters, but with different timing. The median time to the start of the PCR-defined infectious period was 43.9 hpc for unvaccinated sentinels in cluster I and 67.9 hpc for unvaccinated sentinels in cluster III; all sentinels in these groups became positive within approximately 80 and 104 hpc, respectively. Swab loads ranged from 3.4 x 10^2 to 2.2 x 10^5 Geq/ml (Figures 2 and 3; Supplementary Tables 2 and 3). In cluster II, which contained vaccinated sentinels, two birds were already positive at 24 hpc and the last became positive at 104 hpc; the estimated median was 55.9 hpc. Numbers of PCR-positive sentinels by day and replicate are shown in Figure 2, and the corresponding Cq values are provided in Supplementary Table 2.

**Figure 3.**
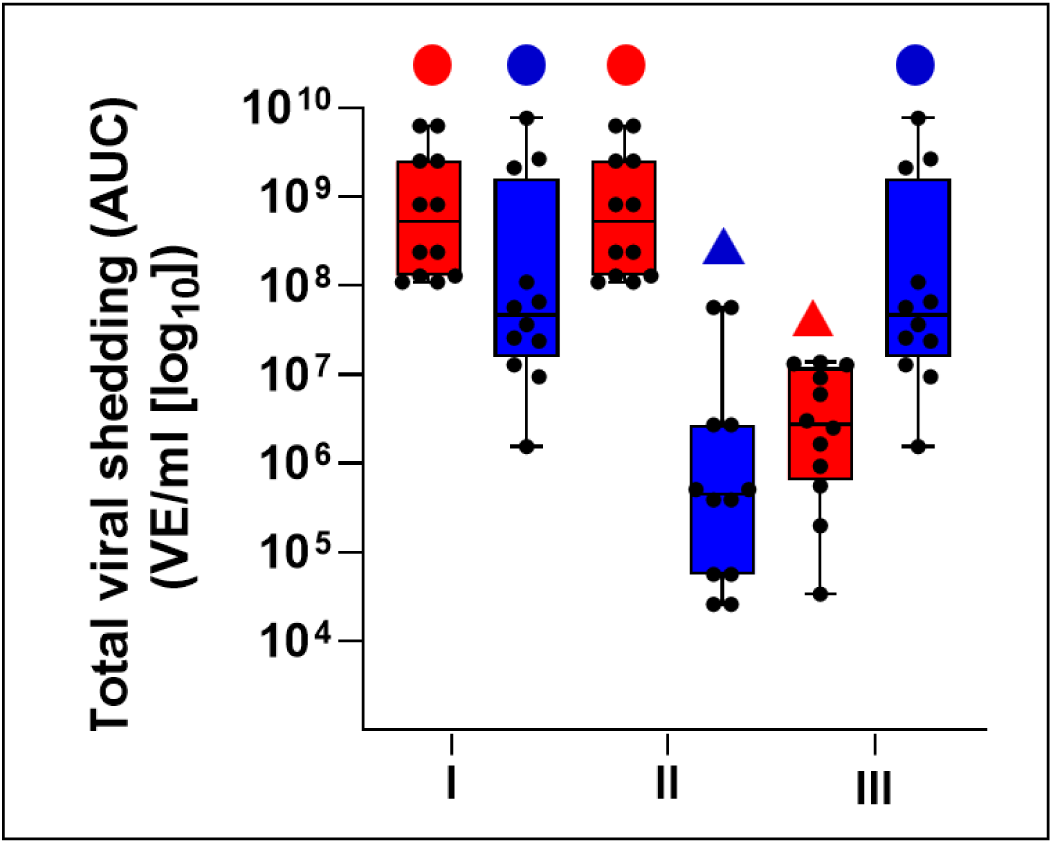
Total viral shedding (AUC) in vaccinated and unvaccinated animals. Total viral shedding per animal was determined as the area under the curve (AUC) of virus equivalents (Geq/ml, log₁₀) over time. Data are shown separately for clusters I–III. Box plots represent the median, interquartile range, and min– max values, with individual animals shown as symbols. Colors indicate the mode of infection (red, inoculated animals; blue, sentinel animals), while symbols indicate vaccination status (dots, unvaccinated; triangles, vaccinated). Cluster I represents the unvaccinated control setting, whereas clusters II and III illustrate viral shedding following transmission between vaccinated and unvaccinated animals.

#### Vaccination markedly reduces cumulative viral RNA shedding

Cumulative oropharyngeal viral RNA shedding was summarized as the area under the curve (AUC) of genome equivalents over time (Figure 3). Vaccinated infected geese showed markedly lower AUC values than unvaccinated infected geese despite remaining in the experiment for longer. Interpretation of AUC values in unvaccinated birds requires caution because many were euthanized or removed early for animal-welfare reasons. In particular, the comparatively low AUC values of unvaccinated sentinels in cluster I reflect removal soon after they met the RT-qPCR criterion for infectiousness and before more extensive shedding could develop.

### Vaccination reduces H5N1 transmission but does not eliminate it

Rapid secondary transmission was a potential source of bias, particularly in clusters containing unvaccinated sentinels. Even though sentinel geese were removed as early as possible after testing positive for virus shedding, infected naive sentinel geese could shed large amounts of viral RNA before and could have infected other sentinels before removal. To address this feature of the experiment, we estimated transmission under two complementary assumptions. The seeder-only model treated only inoculated infectious geese as sources of infection, whereas the all-infectious model also allowed newly infected sentinels to contribute to transmission until removal. The first model reflects the intended seeder-to-sentinel transmission contrast; the second captures the possibility of secondary transmission within each experimental group. Detailed model definitions are provided in Materials and Methods.

#### Vaccinated geese have a longer PCR-defined infectious period but lower transmission rates

The estimated PCR-defined infectious period was longer in vaccinated than in unvaccinated geese (4.24 days [95% CI, 1.81-6.37] versus 2.91 days [1.29-4.41]; p=0.007). Within vaccination strata, infectious-period estimates did not differ significantly between inoculated and contact-infected animals. Because infectiousness was defined from serial RT-qPCR results, this duration represents persistence of the molecular shedding signal used by the transmission model rather than direct measurement of viable-virus shedding.

Transmission-rate estimates were similar under the seeder-only and all-infectious assumptions (Table 1), indicating that the main conclusions were not strongly dependent on whether secondary transmission from sentinels was included. Relative to transmission among unvaccinated geese, vaccination reduced the estimated transmission rate both when vaccinated animals were recipients and when vaccinated animals were infectious donors, although these individual contrasts were not statistically significant (p>0.05). Under the all-infectious model, transmission between vaccinated animals was significantly lower than transmission between unvaccinated animals (p=0.03), corresponding to an estimated 70.5% reduction in the transmission rate.

**Table 1.** Estimated parameters for transmission rates (β) and infectious periods (T).

| Transmission Pathway<sup>a</sup> | Infectious Period ($T$) | Transmission Rate ($\beta$)<sup>b</sup> | |
| --- | --- | --- | --- |
|  |  | Homogeneous | Heterogeneous |
| Unv → Unv (cluster I) | 2.91 (1.29-4.41) | 1.61 (0.85-2.73) | 1.63 (0.96-2.74) |
| Unv → Vac (cluster II) | 2.91 (1.29-4.41) | 0.83 (0.44-1.4) | 0.9 (0.53-1.55) |
| Vac → Unv (cluster III) | 4.24 (1.81-6.37) | 0.95 (0.51-1.61) | 0.87 (0.46-1.61) |
| Vac → Vac | 4.24 (1.81-6.37) | - | 0.48 (0.2-1.16) |

<sup>a</sup> This indicates the direction of transmission from vaccinated (Vac) or unvaccinated (Unv) seeders to Vac or Unv sentinels.
<sup>b</sup> Transmission rates are presented for the scenarios considered. Homogenous groups (virus transmission considered from seeders only) and heterogenous groups (virus transmission considered from all infectious (vaccinated and not vaccinated) animals). Note that in the Homogeneous scenario $\beta$ from vaccinated seeders to vaccinated sentinels can not be estimated.
Values in parentheses are the 95% confidence intervals.

#### Experimental reproduction numbers separate vaccine effects on susceptibility and infectiousness

Because the two analytical approaches yielded similar transmission-rate estimates (Table 1), their reproduction-number estimates were also closely aligned (Figure 4A). We therefore focus on the all-infectious model. The baseline reproduction number among unvaccinated geese was R_0_=4.7 (95% CI, 2.8-8.0). Transmission from unvaccinated infected geese to vaccinated recipients was lower (R_s_=2.6 [1.6-4.5]), as was transmission from vaccinated infected geese to unvaccinated recipients (R_i_=3.7 [2.0-6.8]). The derived reproduction number for transmission in a fully vaccinated population was R_vacc_=2.0 (0.8-4.9). Thus, vaccination reduced the point estimate substantially, but the estimate remained above one and its confidence interval included one. The corresponding vaccine-efficacy estimates separated the effects on susceptibility and infectiousness and yielded a combined transmission effect of approximately 57% (Figure 4B).

**Figure 4.**
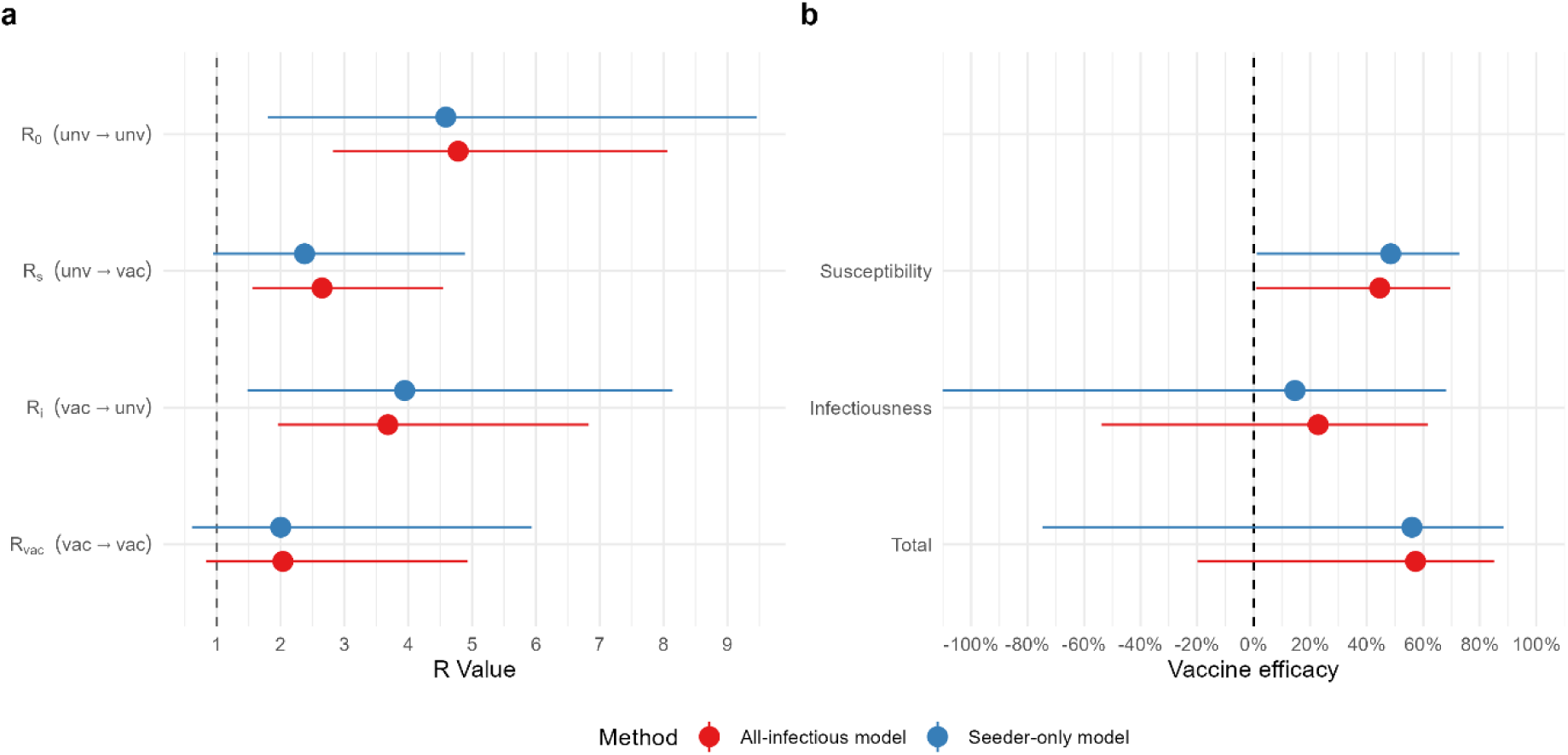
Forest plots of reproduction numbers (A) and vaccine-efficacy estimates (B). Points show mean estimates and bars show 95% confidence intervals for the two analytical assumptions: the seeder-only model (blue), which attributes transmission only to inoculated infectious animals, and the all- infectious model (red), which also allows secondary transmission from infected sentinels. R_0_ denotes transmission among unvaccinated geese; R_s_, transmission from unvaccinated infected geese to vaccinated recipients; R_i_, transmission from vaccinated infected geese to unvaccinated recipients; and R_vacc_, transmission in a fully vaccinated population. Arrows indicate the direction of transmission: vac, vaccinated; unv, unvaccinated.

## Discussion

This study shows that small, controlled transmission experiments can resolve vaccine effects that are not captured by clinical protection or viral shedding alone. Vaccination completely prevented clinical disease in directly challenged as well as in sentinel geese, yet all vaccinated seeders became RT-qPCR-positive and transmission became evident. Relative to the baseline estimate among unvaccinated geese (R_0_=4.7, cluster I), transmission was reduced both to vaccinated recipients (R_s_=2.6, cluster II) and from vaccinated breakthrough-infected donors (R_i_=3.7, cluster III), yielding a derived R_vacc_ of 2.0 for a fully vaccinated population. Under the intensive exposure conditions used here, vaccination therefore reduced transmission substantially but did not reduce the point estimate below the threshold of one.

The subdivision into susceptibility and infectiousness is biologically informative because these components have different implications for control. A vaccine that mainly reduces susceptibility lowers the chance that exposure establishes infection, whereas a vaccine that mainly reduces infectiousness limits onward spread once breakthrough infection has occurred. In the present experiment, a limited vaccine effect on susceptibility (45%) and an apparent even lower effect on infectiousness (23%) were observed. Consequently, the clinical protection observed in vaccinated geese is no indication for sterile immunity or as evidence that breakthrough cases are epidemiologically negligible. This distinction is particularly important for HPAIV vaccination programs in which clinically protected infected birds may be more difficult to detect without active virological surveillance [5,11]. On the other hand, all vaccinated animals that became infected, either after direct inoculation or by exposure to seeder birds, cleared the infection by the end of the experiment, suggesting effective immune-mediated clearance. This should be considered when alternative HPAIV control strategies that include vaccination campaigns are envisioned.

A second important finding was the mismatch between viral RNA shedding and observed transmission. Vaccinated infected geese had markedly lower cumulative oropharyngeal RNA shedding, yet the reduction in R_i_ was comparatively modest. At the same time, vaccinated geese had a longer PCR-defined infectious period, most likely because they survived and remained under observation while low-level RNA could still be detected. RT-qPCR is therefore highly useful for defining infection histories, but persistence and quantity of viral RNA are not equivalent to viable-virus shedding or to infectiousness. For this host-pathogen-vaccine combination, the results caution against using reductions in molecular shedding as a quantitative surrogate for reductions in transmission without validation against transmission data [12].

The two analytical approaches produced similar estimates despite different assumptions about secondary transmission. This agreement is reassuring because rapid transmission from newly infected naive sentinels was a realistic concern in the small groups. The seeder-only model most directly represents the intended experimental contrast, whereas the all-infectious model better reflects the complete transmission process during the interval before infected sentinels were removed. Agreement between the approaches suggests that the estimated vaccine effects were not driven primarily by the handling of secondary transmission in this experiment.

The reproduction numbers reported here should nevertheless be interpreted as experimental transmission parameters, not as direct estimates of flock-level reproduction numbers in the field. Contact structure, group size, husbandry, host heterogeneity, environmental persistence of virus, and exposure intensity are deliberately constrained in a challenge experiment. The high inoculation dose (10^6 EID50) may also have reduced the ability to resolve partial protection against acquisition of infection, because all directly challenged vaccinated geese became infected. Conversely, early removal of unvaccinated infected birds for animal-welfare reasons shortened observation of their natural shedding course. These features can affect absolute R estimates in different directions and argue against interpreting the experimental values as simple upper or lower bounds for field transmission. Yet, this approach will facilitate comparisons of vaccine efficacy among different vaccines investigated in this model system. Several modelling assumptions also affect inference. Within-group heterogeneity in immune responses [23] and contact patterns is not represented explicitly, and the complementary log- log models approximate transmission hazards over discrete observation intervals. The relatively small number of replicated groups necessarily produces wide confidence intervals, most notably for R_vacc_. These limitations do not invalidate the comparative framework, but they emphasize that point estimates should be interpreted together with their uncertainty.

The experimental design was intentionally constrained by animal-welfare considerations. Replicated groups of three seeders and three sentinels provided the contrasts required to estimate susceptibility and infectiousness effects without adding a separate fully vaccinated challenge-transmission arm. This is relevant for severe challenge models such as HPAI, in which increasing sample size can carry substantial welfare costs [13]. Future studies could use simulation-based power calculations tailored to the transmission model and could evaluate whether fewer or differently balanced groups retain adequate precision.

For broader application, two refinements appear particularly important. First, infectiousness should ideally be informed by isolation or titration of viable virus in addition to RT-qPCR, allowing the molecularly defined infectious period to be calibrated against true transmission competence. Second, challenge-dose titration could identify exposure levels (e.g. ID_50_) that remain reproducible while better preserving gradations in vaccine-induced protection. Both refinements would strengthen comparisons between vaccines, virus strains, host species, and experimental laboratories.

Taken together, these results support experimental estimation of reproduction numbers as a complementary endpoint for vaccine evaluation. The approach directly quantifies the population-level consequence of vaccination, separates effects on acquisition and onward transmission, and exposes discrepancies between surrogate virological endpoints and transmission itself. For HPAIV H5N1 in geese, the tested vaccine provided complete clinical protection and reduced transmission, but breakthrough infection and onward spread remained possible under intense experimental exposure. Incorporating direct transmission endpoints alongside safety, immunogenicity, clinical protection, and shedding should therefore provide a more informative basis for comparing vaccines intended to contribute to pathogen control.

## Materials and Methods

We used a controlled HPAIV H5N1 transmission experiment in domestic geese to estimate how vaccination altered susceptibility to infection and infectiousness after breakthrough infection. Geese were selected as a relevant poultry host because commercial production commonly involves outdoor access, increasing opportunities for exposure to avian influenza viruses [14].

### Animal experiments

Goslings of the same hatch-day of a commercial breed of German laying geese were obtained from a commercial hatchery and housed as described previously [15]. After a three-week comprehensive acclimatization phase, animals were randomly separated into two groups, consisting of 24 vaccinees and 24 non-vaccinees. An additional 24 unvaccinated geese were retained as a control group. During the experiment, animals were under continuous, video-assisted supervision by veterinary staff and animal caretakers, realized by at least two daily controls in person and continuous video supervision, supported by a remote-controlled light monitoring regime. Occurrence of any onset of clinical disorders prompted veterinary diagnosis and, if unrelated to HPAI, a treatment chain as described [15].

### Ethics statement

All animal experiments have been granted permission by an independent ethics committee of the German Federal State of Mecklenburg-Western Pomerania (LALLF 7221.3-1-037/23-1). The animal trials took place in the biosafety level 3 (BSL3) facilities of the Friedrich-Loeffler-Institute (FLI) on the Isle of Riems, Greifswald.

### Vaccine

An RNA-based vaccine was used for immunization; this vaccine had not been licensed for use in Europe at the time of testing. This is why the producer provided vaccine doses on the basis of MTA/NDAs for exclusive experimental use at FLI.

### Experimental Design

At three and seven weeks of age, geese were immunized twice with the Respons vaccine as indicated in Table 2; Figure 5. Three weeks after the second vaccination, a challenge trial was conducted and geese were randomly allocated to three distinct experimental clusters as outlined in Figure 2: In **Cluster I**, four subclusters of three unvaccinated “seeder”-geese each were inoculated oculo-nasally with 0.5 ml containing 10^6^ EID_50_ of the H5N1 clade 2.3.4.4b HPAIV isolate *A/chicken/DE-NI/AI 4286/2022*, a genotype Eurasian (EA) AB strain of HP phenotype in waterfowl [15,16]. Following an incubation period of 16 hours, each subcluster of three geese was co-housed with three unvaccinated, non-inoculated “sentinel”-geese. Analogous clusters were formed with unvaccinated, inoculated “seeder”-geese and vaccinated “sentinel” geese (**Cluster II**) and with vaccinated, inoculated “seeder” geese and unvaccinated “sentinels” (**Cluster III**). Two of the four groups in each cluster were housed in the same stable but spatially separated, with each group having its own drinking water and feed supplies, resting area, and bathing area.Altogether, 72 geese were used and allocated to six high containment stables (BSL3).

**Figure 5.**
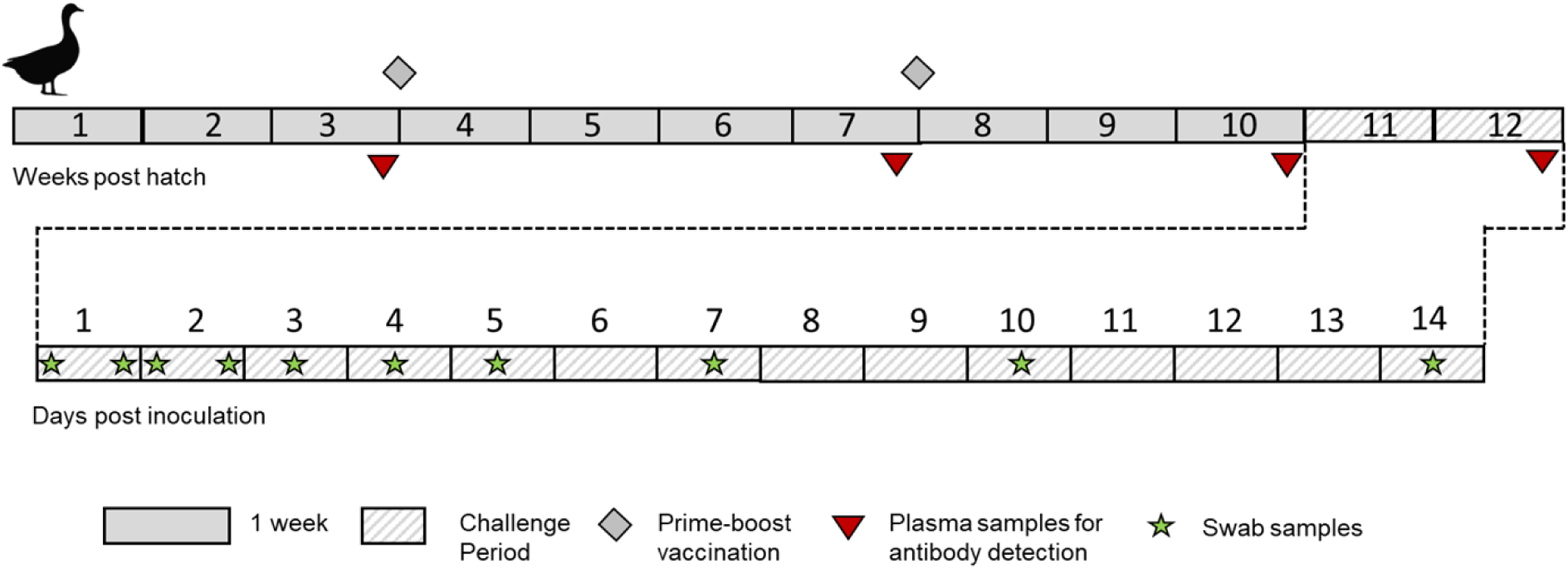
Vaccination and sampling scheme of geese.

**Table 2.** Characteristics of vaccine used.

| Vaccine producer | Vaccine | Batch number | Type [gs/GD clade] | Dose and application |  |
| --- | --- | --- | --- | --- | --- |
| CEVA | RESPONS AI H5 | Lot 0409LF | Amplicon (H5) [2.3.4.4b] | 0.2 ml | i.m., caudal femoral muscle |

Animals were examined during the whole challenge period both physically and via a recording video system: twice daily during the acute phase of infection. Defined criteria for clinical scoring were applied as described [15], that included human endpoints for euthanasia of severely diseased geese. In addition to clinical criteria, unvaccinated sentinel geese (Clusters I and III) were removed once recognized by RT-qPCR (Cq ≤ 35.99) to shed virus and before severe clinical signs could develop. The vaccinated animals, regardless of whether belonging to the seeder (Cluster III) or sentinel group (Cluster II), did not develop any clinical signs and therefore remained in the group for the entire observation period of 14 days. For determination of virus shedding, separate oropharyngeal as well as cloacal swab samples were taken from all animals at indicated hours after inoculation (hpi), i.e twice on day one and two (16, 24, 40 and 48 hpi), and once daily on days 3 (64 hpi), 4 (88 hpi), 5 (112 hpi), 7 (160 hpi), 10 (232 hpi) and 14 (328 hpi). Staff entered BSL3 stable units in overpressurized suits and proceeded with manipulations in the direction from Cluster III to Cluster I units. Between each cluster, suits and attached boots were cleaned (water) and disinfected using Virkon S (1%) in a separate room.

### Real-time RT-PCR (RT-qPCR)

For RNA extraction, swab samples and tissues were treated as described previously [15,16] and tested for presence of viral RNA in a previously published RT-qPCR protocol detecting a conserved target in the M gene segment [17]. Cq values were semi-quantitatively correlated with virus genome equivalents (Geq) based on an intra-assay calibration curve of a defined HPAIV H5N1 virus stock with known TCID_50_ [16]. For the purposes of R assessment, a Cq value of ≤ 35.99 was considered AIV RNA-positive (corresponding to 400 virus genome equivalents/ml), and the respective animal was designated as “AIV infected”. In the event that the Cq value remained ≤ 35.99 in the ensuing measurement interval, i.e., at the same day during day 1 and 2 or on a minimum of two consecutive days after day 3, the animal was designated as “infectious” and, hence, categorized as a “shedder”.

Total virus shedding per animal was quantified by calculating the area under the curve (AUC) from precalculated virus equivalents (Geq/ml). For each animal, daily virus shedding was estimated as the mean of the Geq/mL values from the current and subsequent sampling days, multiplied by the time interval between the two sampling points (in hours). The AUC values obtained for each interval were then summed to yield the total virus shedding per animal (see Figure 4 and Supplementary Table 5).

### Serology

Heparinized blood samples were collected as indicated in Figure 5. Heparinized plasma was examined in two commercial ELISA test kits for Influenza A-specific antibodies against the NP or H5 proteins (ID screen Influenza A Antibody Competition Multispecies ELISA; ID screen Influenza competitive subtype-specific H5 kit; ID Screen Influenza H5 Antibody Competition 3.0 Multi-species) according to the instructions of the manufacturer. Additionally, all samples were tested in a haemagglutination inhibition (HI) assay against the homologous H5N1 clade 2.3.4.4b challenge antigen.

### Statistics

#### Sample size determination

The sample size was determined based on previous experimental studies investigating transmission parameters in animal models. In particular, we followed the approach described by [18], who estimated transmission rates in controlled experiments and provided practical guidance for sample size selection in such settings. A simplified power analysis was performed. Assuming that the transmission process can be approximated by a Poisson-like distribution, we estimated the minimal detectable difference in R_0_ under a scenario with twelve seeders and twelve sentinels per cluster.

The final sample size was derived based on the following assumptions: A statistical power of at least 0.8 was targeted. Informed by prior studies, the expected reproduction number under high transmission conditions was assumed to be approximately 10.0, while in groups with reduced transmission due to vaccination, the reproduction number was expected to be around 0.5. A balanced design of seeders and sentinels was favored, as recommended by ([18]), to optimize statistical efficiency and interpretability. Based on these parameters, a study design with three seeders and three sentinels in each replicate and four replicates per transmission scenario was deemed sufficient.

#### Estimation of transmission parameters and reproduction numbers

Animals in the experiment were sampled and tested regularly at defined intervals, providing information for each time interval on whether each individual animal within the experimental groups can be classified as susceptible (S), infected but not yet infectious - commonly defined as latently infected or exposed (E) -, infectious (I) and recovered or dead (R). These classifications of the different states of infection were used to prepare the data as exemplified elsewhere [14,19] for the estimation of the following parameters: 1) the latent period (*L*), which is the length of time between an animal becoming infected and infectious, 2) the transmission rate parameter β, which is the average number of secondary infections an infectious individual generates per unit of time, 3) the infectious period (*T*), which is the length of time an infected individual is contagious and, 4) the Reproduction number (*R*). For data preparation and analyses we considered an animal to be infected *and* infectious when it was RT-qPCR positive for at least two consecutive sampling intervals with Ct values < 35.99 (see section RT-qPCR).

#### Transmission rate ***β***

We first estimated *L* using parametric survival regression models. We then used this information to prepare the data for the estimation of β. Two scenarios were considered for this estimation: (i) we considered that only the groups of vaccinated or unvaccinated inoculated- infected animals could transmit infection to either vaccinated or unvaccinated groups of contacts (homogeneous infectious or susceptible groups, referred to as the”seeder-only model”) and, (ii) in this second scenario we considered that both unvaccinated (unv) and vaccinated (vac) infectious animals can transmit infection within each experimental group (heterogeneous groups, referred to as the “all infectious model”).

Scenario (i) was considered because unvaccinated sentinel animals (clusters I and III) were removed once they showed clinical signs and tested positive by RT-qPCR. We therefore assumed that these animals made no, or only a negligible, contribution to transmission to the remaining susceptible contact animals during the short period they remained in the group. In contrast, under scenario (ii), no such assumption was made, and these animals were considered infectious to the remaining susceptible animals during the interval between infection and removal from the experimental group. The analytical approach under this scenario allows all experimental animals to be retained for the duration of the experiment under conditions where, due to specific experimental constraints, frequent sampling and testing for prompt removal, as performed in this study, would not be feasible.

To estimate ß, we applied a modeling approach based on generalized linear models (GLMs) with a binomial error distribution and a complementary log-log (cloglog) link, following the method proposed by [20] and used in several experimental transmission studies in the veterinary field (i.e., [14,21–23]). In these models, the outcome of interest was the number of new infections (cases) among susceptible sentinels during discrete observation intervals. For a more detailed description of the analysis performed here for both scenarios, we refer to Dekker et al. [22].

##### i. Estimation of β assuming homogeneous groups – seeder-only model

In this scenario we fitted the binomial GLM to the number of new cases (C_t_) among susceptibles (S*_t_*), offset by the logarithm of the product of the infectious proportion and the time interval 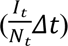 as follow:

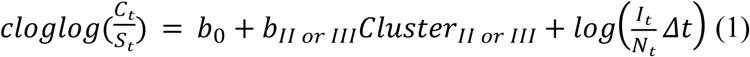

Where *Cluster* is an indicator variable that identifies either Cluster I (unvaccinated (unv) donors - unvaccinated sentinels), Cluster II (unvaccinated (vac) donors - vaccinated sentinels) and, Cluster III (Vaccinated donors - unvaccinated sentinels) (Figure 1) and allows statistical comparison of the transmission rates among the different Clusters. The following *βs* were derived from this model:

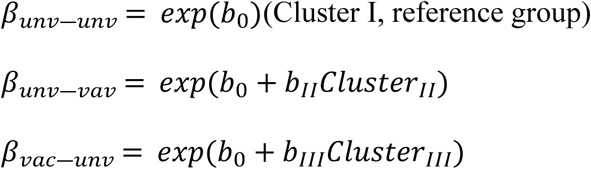

##### ii. Estimation of β assuming heterogenous groups – all infectious model

Different from scenario (i), in this scenario animals within each experimental group can be *I_unv_*_,*t*_,*I_vac_*_,*t*_, *S_unv_*_,*t*_, *S_vac_*_,*t*_,*C_unv_*_,*t*_ *or C_vac_*_,*t*_, consequently the fitted GLM model considers all these different individuals in the group as follow:

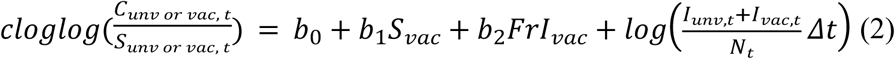

In this model *S_vac_* is an indicator variable equal to 1 when susceptible animals were vaccinated and 0 otherwise; *FrI_vac_* is the fraction of vaccinated Infectious animals. The fitted parameters *b*_1_representing the effect of vaccination on susceptibility and *b*_2_ representing the effect of vaccination on infectiousness when *FrI_vac_* = 1. These parameters were then used to derive the following βs:

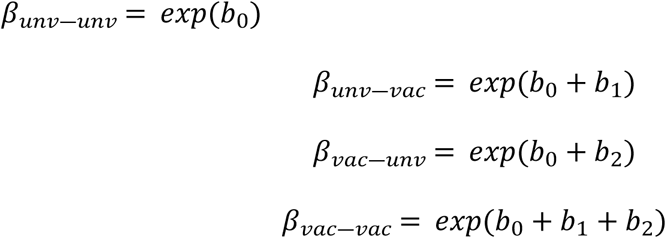

#### Infectious period

The length of the infectious period of vaccinated (*I_vac_*) and unvaccinated (*I_unv_*) infectious individuals was estimated using parametric survival analysis, testing different distributions (Weibull, lognormal and exponential) and comparing their fit based on the models’ Akaike Information Criterion (AIC) values. The Weibull distribution provided the best fit. In these models the vaccination status was introduced as a covariate, to be able to compare whether the infectious period deferred between vaccinated and unvaccinated infectious animals.

#### Reproduction number

The reproduction number R was then calculated as

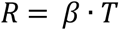

where β is the estimated transmission rate (*β_unv_*_―*unv*_,*β_unv_*_―*vac*_,*β_vac_*_―*unv*_ *or β_vac_*_―*vac*_) and *T*(*^T^unv ^or^*^*T*^*vac*).

When *β_unv_*_―*unv*_ and *T_unv_* are used, we estimate the basic reproduction number R_0._

Analogous estimates were obtained for

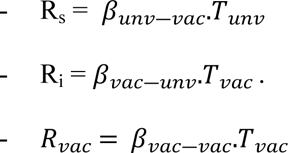

*R_vacc_* was estimated directly when *β_vac_*_―*vac*_ was derived from the GLM for heterogeneous groups. For the scenario where we used the GLM for homogeneous groups, *R_vacc_* was derived after the estimation of vaccine effects as described in the next section.

#### Estimation of vaccine effects

Using the R estimates, we inferred vaccine-induced reductions in susceptibility (*VE_s_*), infectiousness (*VE_i_*) and transmission or total efficacy (*VE_T_*) as follow:

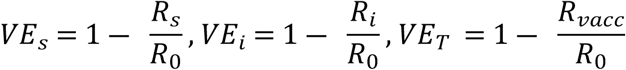

In case of the scenario where homologous groups were assumed, *VE_T_*was calculated as:

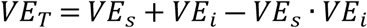

This enabled estimation of the effective reproduction number among vaccinated individuals (R_vacc_), for the scenario considering homogenous groups:

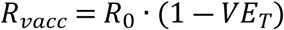

#### Analysis software

All analyses were conducted in RStudio (version 4.3.2) [24], using the packages `purrr` [25]`, here` [26], `tidyverse` [27]`, and MASS` [28]. The library `survival` [29] was used for parametric survival analysis.

#### Data preparation

For data analysis, Cq values obtained from RT-qPCR analysis of oropharyngeal swabs were recorded for each animal and sampling time point and structured on a group-wise basis. Only results from oropharyngeal swabs were considered for R assessment, as these samples showed earlier and more consistent detection of viral RNA and generally lower Cq values, indicative of higher viral RNA loads, compared with cloacal swabs (Supplemental Table 2).

In accordance with the criteria defined above (see chapter RT-qPCR), samples with Cq values ≤35.99 were considered AIV RNA-positive, and the respective animals were classified as AIV *infected*. Animals with Cq values ≤35.99 at the subsequent sampling time point, i.e., during repeated sampling on days 1 and 2 or on at least two consecutive sampling days from day 3 onwards, were classified as *infectious* and categorized as shedders.

For subsequent R assessment, the group-wise Cq value tables were reformatted into .csv files (see data template in Supplemental Materials) and converted into binary matrices. For each animal and sampling time point, Cq values ≤35.99 were coded as 1 (AIV RNA-positive), whereas Cq values ≥36 or samples without detectable viral RNA were coded as 0 (AIV RNA- negative). In cases of intermittent detection, i.e. a single AIV RNA-negative result occurred between two AIV RNA-positive results at three consecutive sampling time points, the intermediate negative result was recoded as 1, and the animal was considered continuously positive over this period.

For animals classified as infectious, the infectious period was defined as the interval from the first AIV RNA-positive oropharyngeal swab associated with the infectious episode to the last AIV RNA-positive oropharyngeal swab before it remained negative or was removed from the experiment. Animals were considered recovered, and therefore no longer infectious, when no further AIV RNA-positive results were detected following the infectious period. For animals withdrawn from the experiment, no further observations were available from the time of withdrawal onwards; these missing observations were therefore distinguished from AIV RNA-negative results and were not interpreted as evidence of recovery.

## Acknowledgments

Ceva Animal Health, Libourne, France, kindly provided the vaccine. We would like to extend our sincere thanks to the laboratory technical team at FLI - Kristin Trippler, Diana Parlow and Cornelia Illing - for their essential support in the laboratory. Additionally, we are deeply grateful to the animal caretakers for their dedicated and careful assistance throughout the animal experiments. Christoph Staubach was helpful in the initial attempts of framing the process mathematically. We are indebted to Christophe Cazaban, CEVA, for critically commenting on the draft manuscript.

## Author Contributions

Conceptualization and methodology by JG, JS, TCH; investigation and data curation by RP, CG; formal analysis by JS, RG, JG; mathematical modelling – JS, MH, JG; writing - original draft preparation by TCH, JS, RP, CG, JG; writing - review and editing by all co-authors; visualization by RG, CG, JG; project administration by CG, TCH; funding acquisition by TCH, CG. All authors have read and agreed to the published version of the manuscript. Generative artificial intelligence technologies were used for editorial and linguistic corrections of the manuscript.

## Data Availability Statement

All curated data underlying the findings reported in this manuscript are included in the Supporting Information.

